# Cannabinoids Activate the Insulin Pathway to Modulate Mobilization of Cholesterol in *C. elegans*

**DOI:** 10.1101/2022.07.20.500754

**Authors:** Bruno Hernandez-Cravero, Sofia Gallino, Jeremy Florman, Cecilia Vranych, Philippe Diaz, Ana Belén Elgoyhen, Mark J. Alkema, Diego de Mendoza

## Abstract

The nematode *Caenorhabditis elegans* requires exogenous cholesterol to survive and its depletion leads to early development arrest. Thus, tight regulation of cholesterol storage and distribution within the organism is critical. Previously, we demonstrated that the endocannabinoid (eCB) 2-arachidonoylglycerol (2-AG) plays a key role in *C. elegans* modulating sterol mobilization, but the mechanism is unknown. Here we show that mutations in the *ocr-2* and *osm-9* genes coding for transient receptors potential V (TRPV) ion channels, dramatically reduces the effect of 2-AG in cholesterol mobilization. Through genetic analysis combined with the rescuing of larval arrest induced by sterol starvation we found that the insulin/IGF-1signaling (IIS) pathway and UNC-31/CAPS, a calcium-activated regulator of neural dense-core vesicles release, are essential for 2-AG-mediated stimulation of cholesterol mobilization. These findings indicate that 2-AG-dependent cholesterol trafficking requires the release of insulin peptides and signaling through the DAF-2 insulin receptor. These results suggest that 2-AG acts as an endogenous modulator of TRPV signal transduction to control intracellular sterol traffic through modulation of the IGF-1 signaling pathway.

**Author summary:** Although cannabis extracts have been used in folklore medicine for centuries, the past few years have seen an increased interest in the medicinal uses of cannabinoids, the bioactive components of the cannabis plant, for treatment of many diseases of the nervous system. However, the human body naturally produces endocannabinoids that are similar to the cannabinoids present in Cannabis sativa. Our goal is to understand how endocannabinoids maintain cholesterol homeostasis in animals, underscoring the importance of cholesterol balance for healthy life. Both cholesterol excess and cholesterol deficiency can have detrimental effects on health, and a myriad of regulatory processes have thus evolved to control the metabolic pathways of sterol metabolism. The nematode *C. elegans* is auxotroph for sterols, that is; contrary to mammals they cannot synthesize sterols, therefore, dietary supply is essential for survival. The aim of our study was to elucidate the mechanism by which endocannabinoids abolish larval arrest of *C. elegans* induced by cholesterol depletion. We discovered that endocannabinoids stimulate the insulin pathway, which affects development, reproduction and life span, to modulate mobilization of cholesterol in *C. elegans*. Our studies have important implications for a better understanding of human pathological conditions associated with impaired cholesterol homeostasis.

## Introduction

Cholesterol is essential for a diverse range of cellular processes, including hormone signaling, fat metabolism, and membrane structure and dynamics. Dysregulation of cholesterol and lipid homeostasis can have a major impact on development and disease [1,2]. Cholesterol deficiency can result in blunted steroid hormone production, reduced serotonin levels, vitamin deficiencies, and increased mortality, whereas cholesterol excess is a risk factor for cardiovascular disease, diabetes, neurodegeneration, and inflammation [3–8]. Thus, understanding cholesterol and lipid homeostasis is critical to illuminate aspects of human health and longevity.

The nematode *Caenorhabditis elegans* requires exogenous cholesterol because cannot synthesize it de novo [9]. In *C. elegans*, cholesterol regulates at least two processes. First, it is required for growth and progression through larval stages, as well as for proper shedding of old cuticles during larval molting events [10]. Second, it regulates entry into a specialized diapause stage adapted for survival under harsh conditions, called the dauer larva [11]. Tight spatial and temporal regulation of uptake, storage, and transport of sterols to appropriate subcellular compartments is required for cholesterol to exert its diverse cellular functions [10].

Despite the pivotal role of sterols in *C. elegans* development, the regulation of cholesterol metabolism is only now beginning to be understood. Previously, it was shown that worms grown in the absence of cholesterol arrest as dauer-like larvae in the second generation [9]. The sterols that govern dauer formation are bile-acid-like hormones called dafachronic acids (DAs) [11]. Molecular mechanisms underlying their function have been intensely studied. DAs inhibit dauer formation by binding to the nuclear hormone receptor DAF-12, which, in the absence of DAs, activates the dauer program [12,13]. Even though cholesterol is associated with cell membranes and interacts with multiple lipid species, very little is known about how lipids influence cholesterol trafficking. It was recently discovered that the *C. elegans* glycolipids, phosphoethanolamine glucosylceramides (PEGCs), stimulates the growth of worms by a yet unknown mechanism under conditions of cholesterol scarcity [14]. We also recently reported that the best studied endocannabinoids (eCBs) 2-arachidonoyl glycerol (2-AG) and arachidonoyl ethanolamine (AEA), which are lipid messengers that elicit a plethora of biological functions in mammals, enhance traffic of cholesterol in *C. elegans* [15]. We found that these eCBs stimulate worm growth under conditions of cholesterol scarcity and reverse the developmental arrest of *ncr-1-2* mutants [15]. *ncr-1* and *ncr-2* encode proteins with homology to the human Niemann-Pick type C (NP-C) disease gene (*NPC-1*) and are involved in the intracellular cholesterol trafficking in *C. elegans* [16]. The mechanism by which these signaling lipids exert their effects on cholesterol homeostasis within large endocrine networks is unknown. Here we show that 2-AG promotes cholesterol mobilization through pathways that are independent of known *C. elegans* cannabinoid-like receptors that mediates regulation of regenerative axon navigation [17] and behavior [18]. We find that mutations in the ocr-2 and osm-9 genes coding for transient receptors potential of the vanilloid subtype (TRPV), ion channels, dramatically reduces the effect of 2-AG in cholesterol mobilization. We also find that the insulin/IGF 1 signaling (IIS) pathway and UNC-31/CAPS, the calcium-activated regulator of dense-core vesicles exocytosis (DCVs), are necessary for 2-AG-mediated stimulation of cholesterol mobilization. This suggests that 2-AG-dependent cholesterol traffic requires signaling of insulin peptides through the DAF-2 insulin receptor. Our results indicate that 2-AG acts as endogenous modulators of TRPV channels to control intracellular sterol traffic through modulation of the insulin/IGF-1 (IIS) signaling pathway.

## Results

### FAAH-4 is involved in cholesterol homeostasis in *C. elegans*

*C. elegans* interrupts reproductive development and arrest as L2-like larvae when grown for two generations without cholesterol [9]. Previously, we have found that this arrest is abolished by supplementation with the eCB 2-AG [15, Fig.1A]. Recent work has revealed that the monoacylglycerol lipase FAAH-4, but not other FAAH enzymes in *C. elegans*, hydrolyzes 2-AG [19]. To test the specificity of eCB signaling we asked whether a *faah-4* deletion (*Δfaah-4*) could enhance the rescuing effect of 2-AG on the arrest induced by cholesterol depletion in *C. elegans*. Indeed, we found that under sterol free conditions exogenous 2-AG (50 μM) significantly increases the formation of adults in *Δfaah-4* animals compared to wild-type animals (Fig 1A). In agreement with this result, *faah-4* animals displayed elevated levels of endogenous 2-AG compared to the wild type (Fig. 1B, [19]). Next, we investigated whether a *faah-4* deficiency relieves the phenotype of mutations that perturb cholesterol transport. We found that FAAH-4 RNAi decreased the Daf-c penetrance in null mutants for the Niemann-Pick homologues *ncr-1-2* (Fig 1C). As expected, FAAH-4 RNAi also enhanced the ability of a low concentration of 2-AG (10 μM) to suppress dauer formation of *ncr-1-2* double mutants (Fig 1C). These results confirm the specificity of 2-AG in cholesterol mobilization and suggest that FAAH-4 is an important enzyme in cholesterol homeostasis.

**Fig 1.**
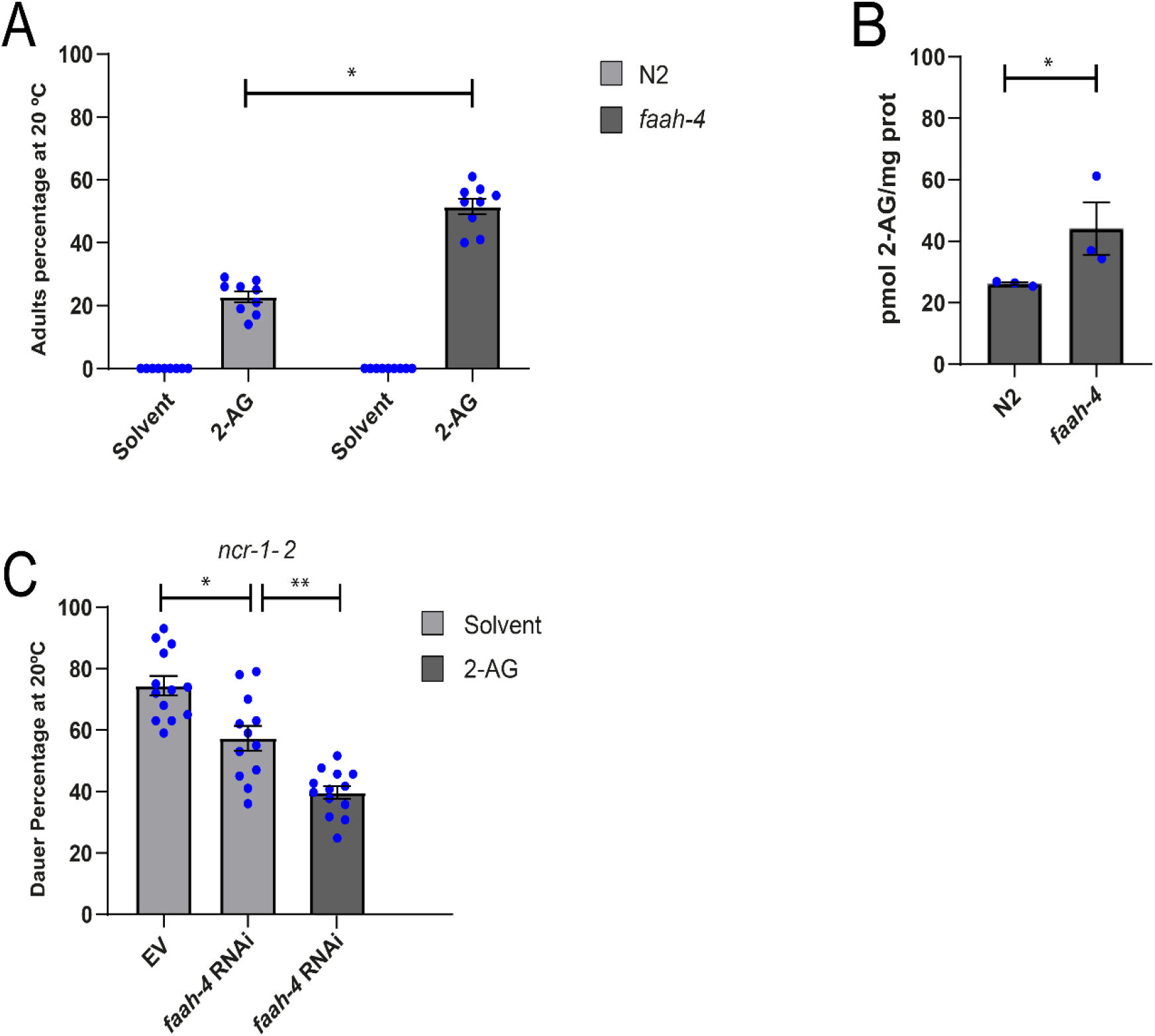
FAAH-4 is involved in cholesterol mobilization by 2-AG. (A) Wild type and *faah-4* animals grown for two generations in the absence of cholesterol at 20 ºC arrest as L2-like larvae. Feeding with 2-AG (50 μg/ml) increases the formation of adults in *faah-4* animals compared with N2 animals. All Pairwise Multiple Comparison Procedures (Dunn’s Method), *p < 0.05. All values from n = 3 independent experiments are shown as Mean ± SEM. (B) Intracellular levels of 2-AG are elevated in *faah-4* animals. T-test, *p < 0.005. All values from n = 3 independent experiments show as Mean ± SEM. (C) *faah-4* RNAi reduces the dauer formation of strain *ncr-1-2* and enhances the ability of low concentrations of 2-AG (10 μM) to suppress the *daf-c* phenotype of these animals. All Pairwise Multiple Comparison Procedures (Holm-Sidak method), *p < 0.002, **p < 0.001. All values from n ≥ 3 independent experiments are shown as Mean ± SEM. N2 is the *C. elegans* wild-type strain.

### 2-AG controls cholesterol homeostasis through NPR-19 and NPR-32 independent pathways

The biological effect of 2-AG in mammals are mediated through its interaction with the G -protein coupled type-1 (CB1) and type-2 (CB2) cannabinoid receptors [20–22]. In a study comparing of vertebrate CB1 to other G protein-coupled receptors (GPCRs) in the *C. elegans* genome, two neuropeptide receptors (NPRs), NPR-19 and NPR-32, were shown to have conservation of the critical amino acid residues involved in eCB ligand binding [17]. NPR-19 has been shown to be a 2-AG receptor that modulates monoaminergic (e.g., serotonin and dopamine) signaling in *C. elegans* [18]. To determine whether NPR-19 is required for the 2-AG-mediated stimulation of cholesterol mobilization, we screened the mutant animals for loss of 2-AG-dependent enhancement of cholesterol trafficking. In particular, we tested the interaction of NPR-19 with the DAF-7/TGF-β pathway, which plays an important role in promoting development by affecting cholesterol metabolism and DA production [14]. *daf-7* mutants constitutively form dauer larvae when grown in normal dietary cholesterol (13 μM) due to impaired cholesterol trafficking [14]. We found that *npr-19* did not increase the penetrance of Daf-C defects seen in *daf-7* mutants at 20°C (Fig 2A). Moreover, dauer formation in both *daf-7* and *daf-7*; *npr-19* mutant animals were rescued by either 2-AG or 2-arachidonoylglycerol-eter (2-AGE) (Fig 2A), a non-hydrolysable analog of 2-AG. In agreement with this prediction, 2-AG relieved the dauer arrest of *npr-19* and npr-32 mutants when cholesterol was depleted from the diet (0 μg/ml) (Fig 2B), indicating that 2-AG stimulation of cholesterol trafficking is independent of NPR-19 and NPR-32. Moreover, 2-AG rescued the arrest of the double mutant *npr-19*; *npr-32* under cholesterol depletion, ruling out the possibility that these two receptors might redundantly act in the eCB effect. Together, these data demonstrate that 2-AG acts in cholesterol mobilization through NPR-19 and NPR-32 independent pathways. We also tested several 2-AG candidates, such as GPCRs and neurotransmitter receptors, identified by examination of protein BLAST data using human CB1 receptors. However, we found that animals mutated in these potential targets were rescued by 2-AG, under cholesterol depletion (S1 Table).

**Fig 2.**
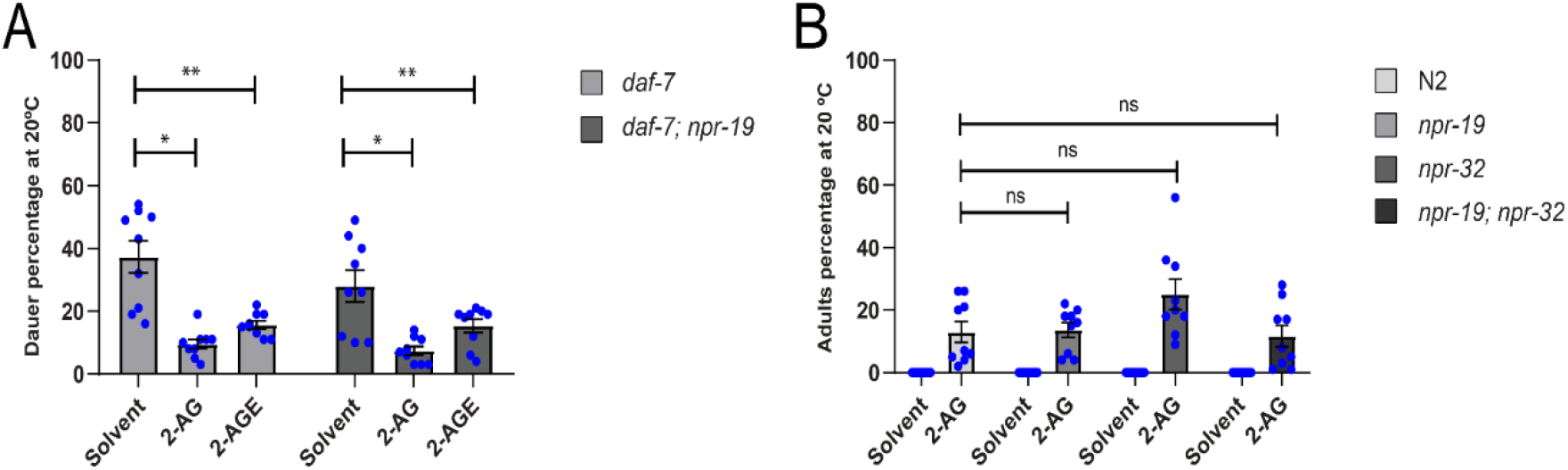
Endocannabinoids stimulate transport of cholesterol by employing pathways independent of *npr-19* and *npr-32*. (A) Endocannabinoid suppress dauer arrest of *daf-7* and *daf-7; npr-19* grown on plates containing 5 μg/ml of cholesterol. Mann-Whitney rank sum test, *p < 0.001, **p < 0.001. All values from n = 3 independent experiments are shown as Mean ± SEM. (B) 2-AG suppress larval arrest of *npr-19, npr-32 and npr-19; npr32* animals grown for two generations on plates containing 0 μg/ml of cholesterol. All values from n = 3 independent experiments show as Mean ± SEM. ns = not significant. N2 is the *C. elegans* wild-type strain.

### TRPV channels are required for 2-AG-dependent cholesterol mobilization

In mammals many cannabinoids can activate transient receptor potential vanilloid (TRPV) channels [23]. *ocr-2* encodes a channel of the TRPV subfamily that functions in *C. elegans* olfaction, nociception and osmosensation [24]. We found that, similarly to *C. elegans fat-3* and *fat-4* mutants, which are deficient in PUFAs and display aberrant cholesterol mobilization [15], *ocr-2* animals displayed a high incidence of arrested larvae in the first generation without cholesterol (Fig 3A and 3B). While 2-AG suppresses dauer formation of *fat* mutants [15], 2-AG was unable to rescue the arrest of *ocr-2* animals starved from cholesterol. To determine the specificity of *ocr-2* requirement for mobilization of cholesterol, we examined mutants for other TRPV channels. The *C. elegans* genome encodes five members of the TRPV family, *ocr-1* through *ocr-4* and *osm-9*. We found that null mutations in *ocr-1, ocr-3* and *ocr-4* produced almost 100% gravid adults in the first generation without cholesterol. In contrast, about 50% of the *osm-9* animals starved of cholesterol exhibited a dauer-like phenotype in the first generation, similar to in *ocr-2* mutants. 2-AG was also unable to prevent the arrest of *osm-9* animals starved from cholesterol (Fig 3A). Together, these data demonstrate that both *osm-9* and *ocr-2* mutations eliminate 2-AG-dependent cholesterol mobilization.

**Fig 3.**
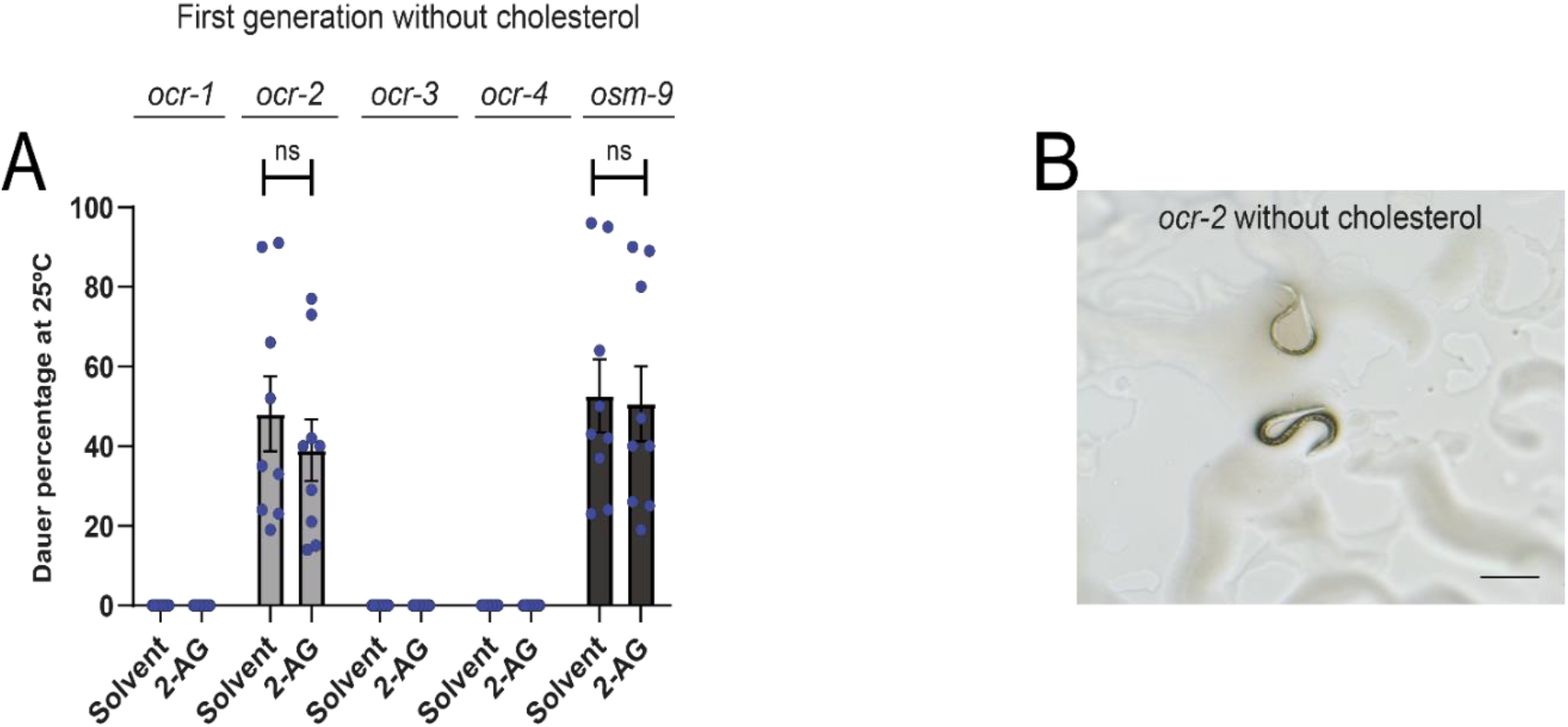
*ocr-2 and osm-9* are required for 2-AG-dependent cholesterol mobilization. (A) Worms were grown in media with 0 μg/ml of cholesterol during one generation at 25 ºC. All values from n = 3 independent experiments show as Mean ± SEM. ns = not significant. (B) *ocr-2* undergoes dauer-like formation in the first generation when grown at 25 ºC in cholesterol-free medium. The black straight line represents 0.25mm.

In *C. elegans*, both, *osm-9* and *ocr-2* TRPV genes are expressed in the ADF serotonergic neurons and also co-expressed in five pairs of non-serotonergic chemosensory neurons: the AWA, ADL, ASH neurons in the head, and the PHA and PHB neurons in the tail [25]. It has been reported that OSM-9 and OCR-2, regulate in ADF, 5-HT biosynthesis through the modulation of the *tph-1* gene expression, which encodes a key enzyme required for 5-HT biosynthesis from tryptophan [26]. A recent report suggested that 2-AG stimulates 5-HT release through a pathway requiring OSM-9, from the serotonergic ADF neurons to modulate *C. elegans* behavior [27]. However, we found that the ability of 2-AG to rescue the development of cholesterol depleted worms is unaffected by mutations in *tph-1* or in mutants in 5-HT receptors (S1 Fig and S1 Table). These results indicate that 2-AG stimulation of cholesterol mobilization in *C. elegans* is independent of 5-HT release.

Both OSM-9 and OCR-2 are thought to function cooperatively and assemble into homo- and hetero-tetramers [28]. Recent electrophysiological studies employing a *Xenopus laevis* oocyte expression system demonstrated that OSM-9/OCR-2 respond to warming, suggesting that these channels cooperatively function as a temperature receptor [29]. Because OSM-9 and OCR-2 channels mediate influx of divalent cations with a preference for calcium [30], we hypothesized that 2-AG is an agonist of channels containing OSM-9 and/or OCR-2 subunits and that activation of the channel is the trigger mechanism for 2-AG-induced mobilization of cholesterol in *C. elegans*. To test this hypothesis, we expressed OSM-9 and OCR-2 in *Xenopus* oocytes and recorded current Two-electrode voltage clamp. Our analysis showed that 2-AG did not elicit currents in *Xenopus* oocytes injected with *osm-9* and *ocr-2* cRNA either alone or in combination (S2 Fig). However, warm stimulus (about 36 ºC) evoked currents in *Xenopus* oocytes simultaneous injected with both *osm-9* and *ocr-2* cRNA (S2 Fig), indicating that the channel was active.

To determine whether 2-AG can induce neuronal activity in vivo in neurons that co-express *osm-9* and *ocr-2* we used calcium imaging. 100 μM 2-AG failed to induce calcium transients in animal that express the GCaMP6 in the ASH neurons (S3 Fig). Since 2-AG did not induce OSM-9/OCR-2 channel activation in vitro or in vivo, it suggests that this compound acts upstream or in parallel of TRPV channels in sensory transduction.

### The cannabinoid based T-Type calcium channel blocker NMP331 antagonizes the action of 2-AG on cholesterol mobilization

Since the effect of 2-AG on cholesterol mobilization appears not to be mediated by CB receptors orthologues (Fig 2), we tested synthetic cannabinoid ligands with the expectation that some of these compounds may act as inverse agonist/antagonist of 2-AG. This approach has been used to identify lipophilic molecules that interact with putative eCB receptors with conserved function but diverge from canonical mammalian receptors [31]. We reasoned that if a synthetic ligand blocks a cannabinoid signaling pathway involved in cholesterol trafficking, the Daf-c defects seen in *daf-7* mutants would be increased. We tested whether a series of CB1/CB2 receptor ligands (NMP compounds) that target both CB receptors and T-type calcium channels [32,33] modulate Daf-c phenotype of *daf-7* (e1372) mutants. Of the four compounds tested, only one compound, NMP331 (named as compound 10 in reference 33), at a concentration of 1μM robustly enhanced the dauer phenotype of *daf-7* at semi-permissive temperature (20°C) (Fig 4A). Supplementation of growth media with an excess of cholesterol lowered dauer formation in *daf-7* exposed to NMP331 (Fig 4B), suggesting that this compound affects sterol mobilization. If NMP331 affects cholesterol mobilization, we predicted that wild-type animals in the presence of this compound would arrest already in the first generation without externally provided sterols. While wild type animals, produced almost 100% gravid adults in the first generation without cholesterol, exposure to NMP331 in the absence of cholesterol resulted in a high incidence (20%) of arrested larvae with typical dauer morphology (Fig 4C and S4A Fig). Furthermore, 2-AG antagonizes the enhancing effect of 1 μM of NMP331 on dauer formation in *daf-7* worms in a dose-response manner (S4B Fig), suggesting that NMP331 acts in the same pathway or in parallel to 2-AG.

**Fig 4.**
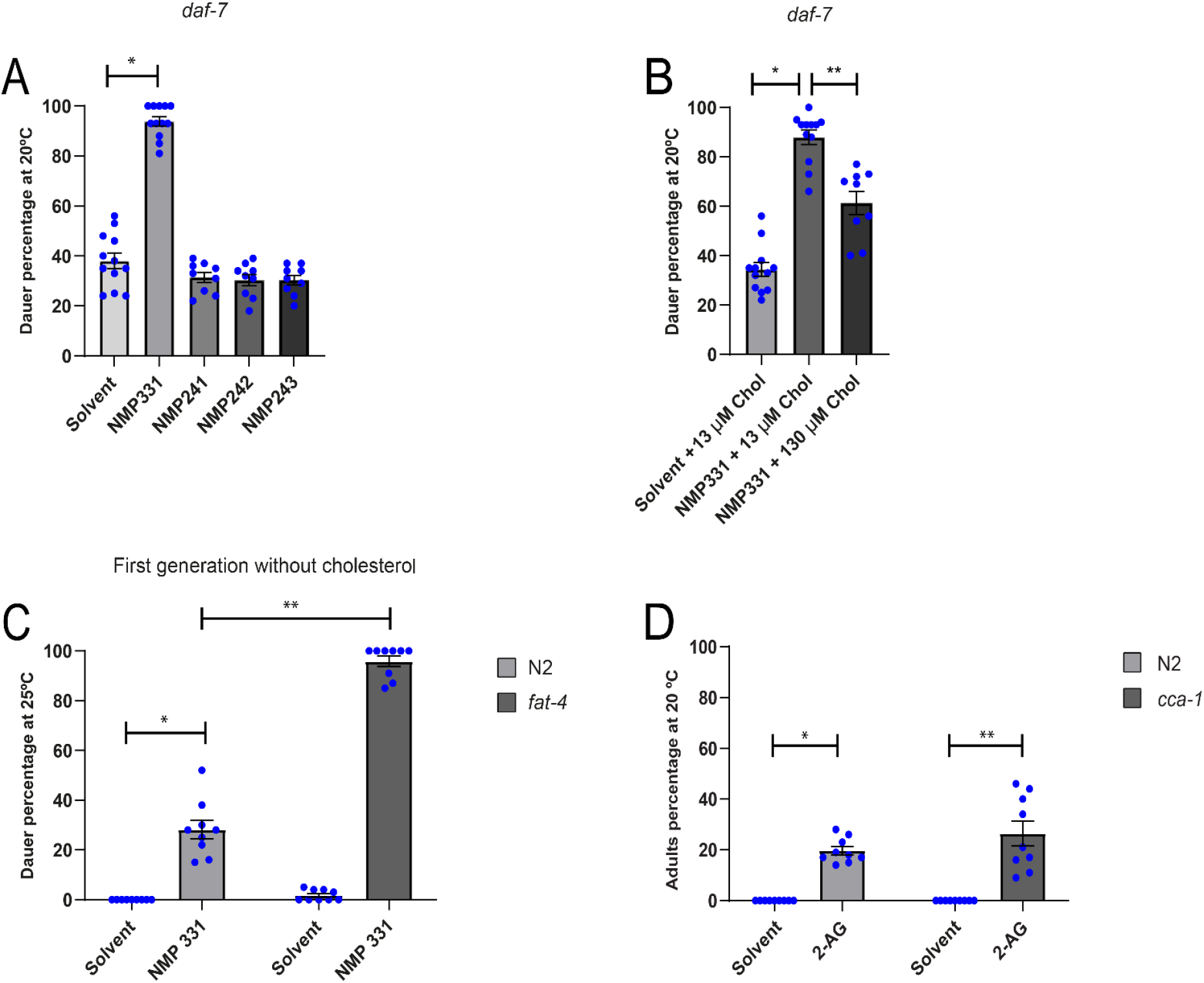
The cannabinoid receptor antagonist NMP331 induces dauer formation. (A) The dauer arrest of *daf-7* worms is augmented by NMP331. All pairwise multiple comparison procedures (Holm-Sidak method), *p < 0.001. All values from n ≥ 3 independent experiments are shown as Mean ± SEM. (B) Dauer arrest induced by NMP331 can be rescued by high cholesterol diet. All Pairwise Multiple Comparison Procedures (Holm-Sidak method), *p < 0.001. **p < 0.001. All values from n ≥ 3 independent experiments show as Mean ± SEM. (C) N2 and *fat-4* animals undergo a dauer arrest induced by 331 in the first generation when grown in cholesterol free medium. Mann-Whitney rank sum test, *p < 0.001. **p < 0.005. All values from n = 3 independent experiments are show as Mean ± SEM. (D) The 2-AG mediated suppression of larval arrest of N2, grown by two generations in medium devoid of cholesterol, is independent of CCA-1. Mann-Whitney rank sum test, *p < 0.001. t-test, **p < 0.001. All values from n = 3 independent experiments show as Mean ± SEM. N2 is the *C. elegans* wild-type strain.

Strikingly, when *fat-4* animals which are unable to synthesize AEA or 2-AG [15] were exposed to NMP331 in a cholesterol depleted medium formed 90% dauers in the first generation, while untreated animals produced mostly gravid adults (Fig 4C).

Using radio-ligand assays and electrophysiology in human embryonic kidney cells, it was determined that compound NMP331 in addition to show high affinity for CB1 receptors is also a blocker of the CaV3.2 T-type calcium channel [33]. Unlike vertebrates that possess three genes that encode T-calcium channels, the genome of *C. elegans* encodes a single T-type channel named *cca-1* [34]. We found that 2-AG-dependent cholesterol mobilization was still present in a *cca-1* mutant (Fig 4D), suggesting that this T-type channel is not the target of NMP331 antagonizing the 2-AG effect in *C. elegans*.

Taken together we conclude that NMP331 impacts cholesterol availability, transport and/or metabolism by antagonizing the stimulatory role of 2-AG in cholesterol mobilization. At this point we do not know whether NMP331 affects other molecular targets, such as ion channels, involved in the cannabinoid-mediated modulation of cholesterol trafficking.

### SBP-1 is required for 2-AG modulation of cholesterol mobilization

To identify potential *C. elegans* genes controlling the regulatory circuit of 2-AG-mediated cholesterol mobilization, we performed RNAi enhancer screen on *daf-7* temperature sensitive mutants, using an RNAi library containing transcriptions factors, transporters and nuclear receptors potentially involved in cholesterol homeostasis (S2 Table). We first surveyed for enhancement of the *daf-7* Daf-c phenotype and, in a secondary survey, screened for loss of 2-AG-dependent rescue of the developmental arrest caused by cholesterol depletion.

We identified two loci, *nhr-8* and *sbp-1*, that in combination with *daf-7* give a strong Daf-c constitutive phenotype (S2 Table). *nhr-8*, codes for a nuclear receptor that plays an important role in cholesterol homeostasis in *C. elegans* [35], while *sbp-1*, encodes the single orthologue of the sterol regulatory element (SREBP) family which regulate transcription of genes required to many aspects of lipid metabolism [36]. We found that supplementation with 2-AG rescued the developmental arrest of *nhr-8* animals depleted of cholesterol (Fig 5A). Thus, *nhr-8* is not positioned within the 2-AG pathway of cholesterol mobilization. In contrast, 2-AG was unable to abolish the developmental arrest induced by cholesterol depletion in *sbp-1(ep79)* animals (Fig 5B). This suggests that 2-AG promotes mobilization of cholesterol, through a pathway requiring the transcriptional activity of SBP-1/SREBP. Consistent with this result, 2-AG was unable to suppress dauer formation in *ncr-1-2* mutants exposed to *sbp-1* RNAi, even though this Daf-c phenotype was largely suppressed by raising the cholesterol concentration in the growth media (Fig 5C). Surprisingly, this high cholesterol concentration did not rescue the Daf-c phenotype of *ncr-1-2* mutants subjected to *sbp-1* RNAi. This suggests that *sbp-1* genetically interacts with the Niemann Pick proteins to regulate intracellular cholesterol trafficking. Finally, we found that *sbp-1* animals displayed elevated 2-AG levels compared with control animals (Fig 5D). This could be due to a compensatory mechanism to optimize cholesterol trafficking in animals depleted of SBP-1.

**Fig 5.**
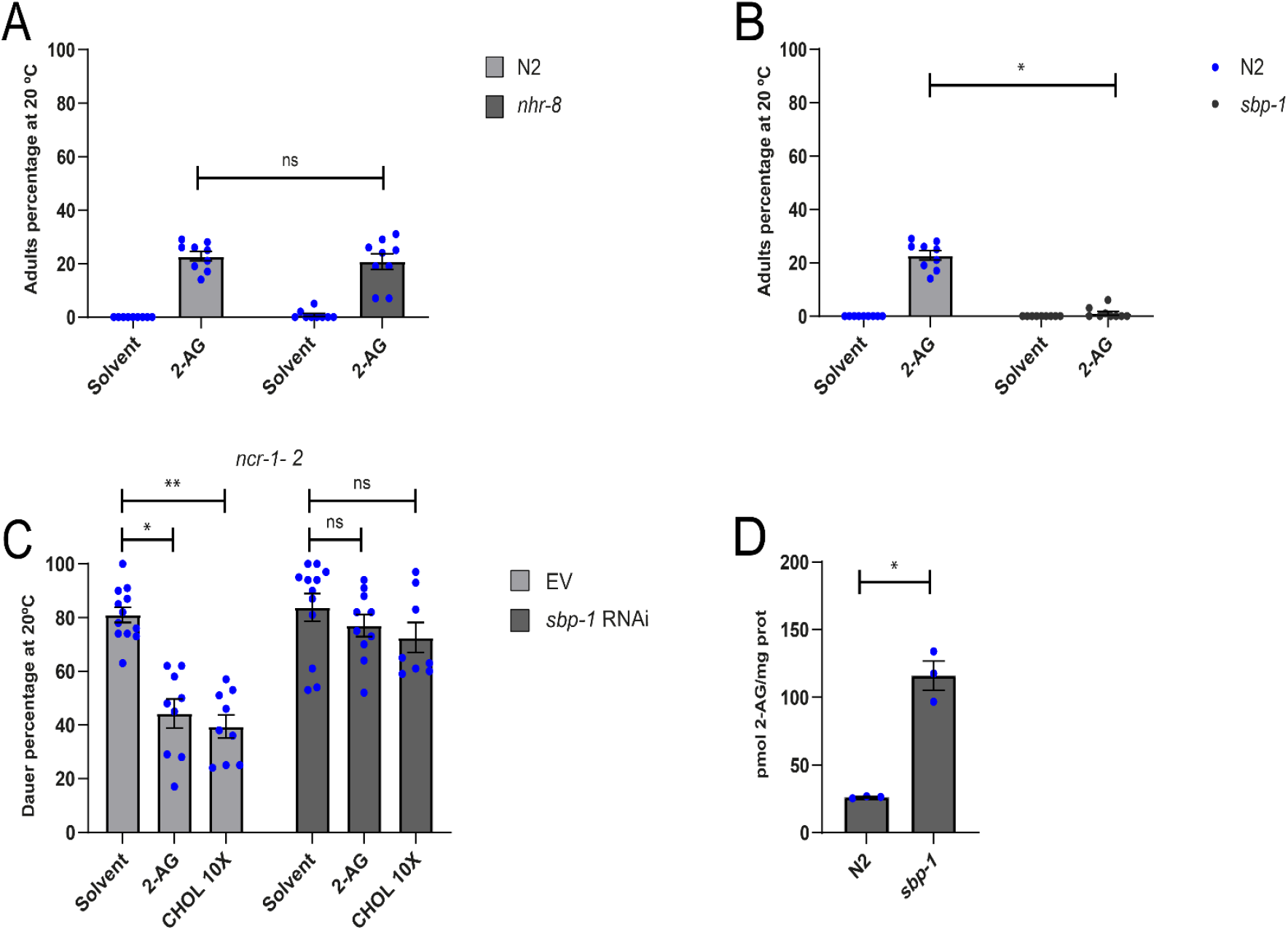
SBP-1 is required for 2-AG modulation of cholesterol mobilization. (A) 2-AG rescue the larval arrest of *nhr-8* worms grown two generations under 0 μg/ul of cholesterol at 20 ºC. All values from n = 3 independent experiments show as Mean ± SEM. ns = not significant. (B) 2-AG is unable to rescue the larval arrest of *sbp-1* grown two generations under 0 μg/ml of cholesterol at 20 ºC. All Pairwise Multiple Comparison Procedures (Dunn’s Method), *p < 0.001. All values from n = 3 independent experiments are show as Mean ± SEM. (C) 2-AG is unable to suppress dauer formation of *a ncr-1-2* strain exposed to *sbp-1* RNAi. Kruskal-Wallis One-way analysis of variance on ranks, *p < 0.001, **p < 0.001. All values from n ≥ 3 independent experiments are show as Mean ± SEM. ns = not significant. (D) 2-AG levels are elevated in *sbp-1* animals. t-test, *p < 0.005. All values from n = 3 independent experiments are show as Mean ± SEM. N2 is the C. elegans wild-type strain.

Taken together, our results suggest that SBP-1 in concert with the Niemann-Pick homologs plays an important role in the 2-AG signal transduction pathway to mobilize cholesterol.

### Mobilization of sterols by 2-AG is controlled by the insulin pathway

The DAF-2/IIS and DAF-7/TGF-β signaling comprise the major endocrine pathways modulating the conversion of cholesterol into DA [13]. In a previous study it was shown that temperature-sensitive *daf-7* mutants are hypersensitive to cholesterol depletion and form dauer larvae in the absence of external cholesterol already at 20°C [14]. More recently, we reported that upon cholesterol deprivation 2-AG rescues the dauer arrest of *daf-7* animals [15]. We determined that 2-AG also prevents dauer formation in mutants which are defective in core components of the DAF-7/TGF-β signaling pathway (S5A Fig), suggesting that 2-AG functions independently of this pathway.

Interestingly, *daf-2(e-1370)* mutants with reduced Insulin-IGF-1 receptor signaling also form about 90% dauers at 20°C in the first generation in the absence of cholesterol (S. Penkov and T. Kurzchalia, unpublished results). Nevertheless, unlike *daf-7* mutants, 2-AG could not suppress the dauer arrest of *daf-2* animals starved from cholesterol a 20°C (Fig 6A). This suggests that 2-AG-mediated mobilization of cholesterol depends on the IIS pathway. We also tested the requirement of phosphoinositide-3 kinase AGE-1/PI3K or the serine threonine kinase AKT-1 that act downstream of the DAF-2 insulin receptor. 2-AG was unable to suppress the dauer formation of both single mutants growing under sterol depleted conditions (Fig 6B-C). Finally, we found that the 2-AG antagonist, NMP331, does not enhance the Daf-c phenotype at 20°C of *daf-2(e1370*) mutants (S5B Fig). This result show that NMP331 requires a functional IIS signaling pathway for its effect on cholesterol trafficking. Taken together, our results suggest that IIS and 2-AG converge in a process essential for cholesterol mobilization.

**Fig 6.**
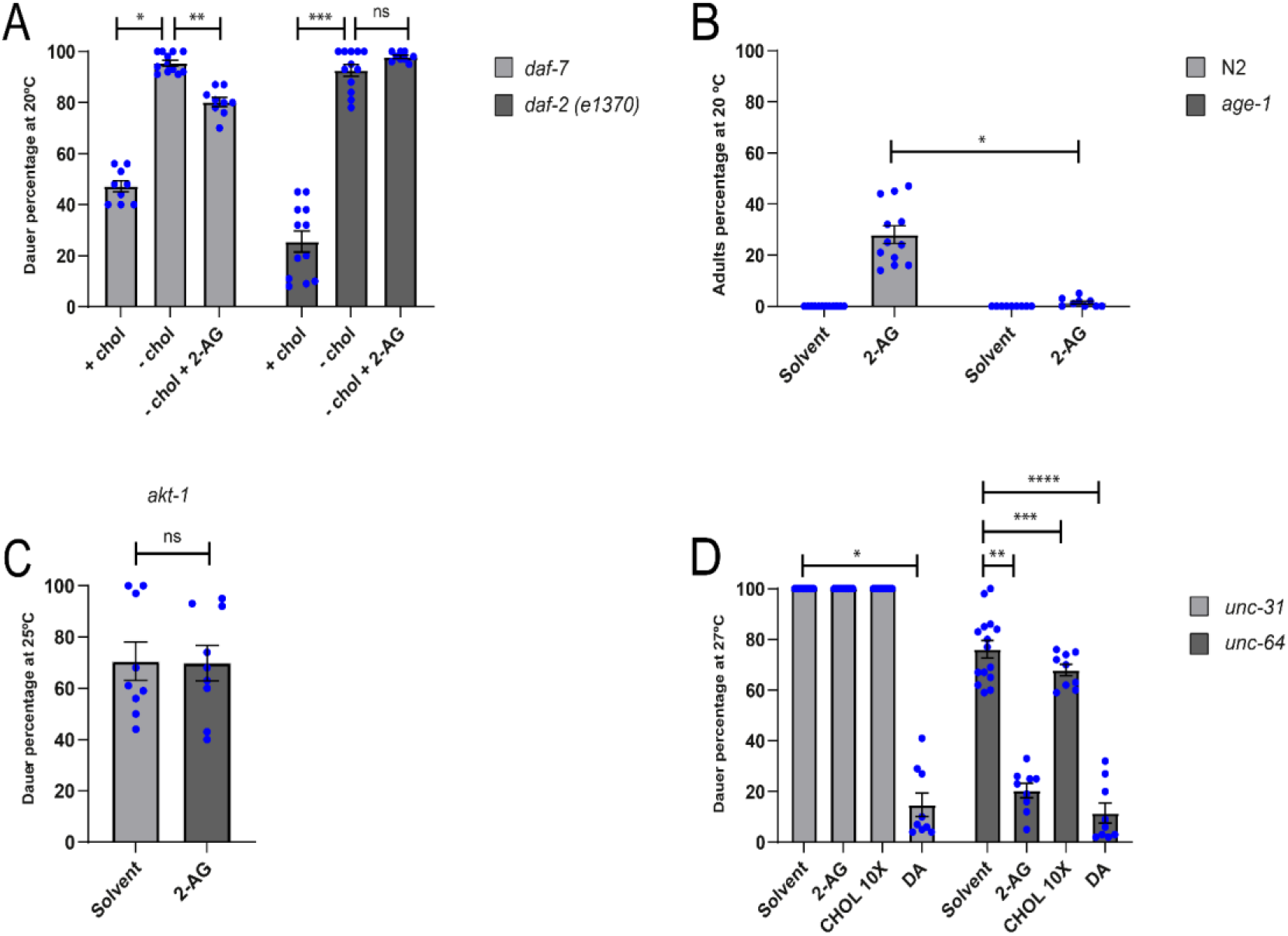
DAF-2 and UNC-31 are required for 2-AG-dependent cholesterol mobilization. (A) *daf-7* and *daf-2(e1370*) were grow at 20 ºC either on cholesterol or in a sterol-free media during one generation. When indicated the media was supplemented with either cholesterol 13 μM or 2-AG 50 μM. All Pairwise Multiple Comparison Procedures (Dunn’s Method), *p < 0.05. **p < 0.05. ***p < 0.05. All values from n ≥ 3 independent experiments show as Mean ± SEM. ns = not significant. (B) N2 and *age-1* were grown for two generations in media with 0 μg/ml cholesterol at 20ºC. Mann-Whitney rank sum test, *p < 0.001. All values from n ≥ 3 independent experiments show as Mean ± SEM. (C) *akt-1* was grown in media with 0 μg/ml cholesterol during one generation at 25 ºC. All values from n = 3 independent experiments show as Mean ± SEM. ns = not significant. (D) 2-AG is unable rescue the daf-c phenotype of *unc-31* grown at 27 ºC. All Pairwise Multiple Comparison Procedures (Dunn’s Method), *p < 0.05. All Pairwise Multiple Comparison Procedures (Holm-Sidak method), **p < 0.05. ***p < 0.05. ****p < 0.05. All values from n ≥ 3 independent experiments show as Mean ± SEM. N2 is the *C. elegans* wild-type strain.

### UNC-31 and HID-1 are required for 2-AG-dependent cholesterol mobilization

We next sought to determine whether 2-AG rescued dauer formation in worms deficient in proteins that act upstream DAF-2, such as UNC-64/syntaxin [37] and UNC-31/CAPS [38]. *unc-64* and *unc-31* mutants have constitutive dauer formation at 27°C but not 25°C (Fig 6D) [39]. Dauer formation of *unc-64* mutant animals was markedly suppressed at 27°C by either 2-AG or DA under normally dietary cholesterol. As expected, high concentrations of cholesterol also suppressed dauer formation of *unc-64* mutants at 27°C (Fig 6D). In contrast, 2-AG was unable to suppress the Daf-c phenotype of *unc-31* mutants, suggesting that its gene product is essential for the 2-AG-dependent mobilization of cholesterol (Fig 6D). Consistent with a role of UNC-31 in cholesterol homeostasis, we found that DA rescues the dauer arrest of *unc-31* animals (Fig 6D). *unc-31* encodes the *C. elegans* homolog of mammalian Ca^2+^ activated protein for secretion (CAPS), required for the regulated release of dense core vesicles (DCVs), which contain biogenic amines, neuropeptide, and insulins [38,40–42]. In addition, 2-AG was unable to rescue the Daf-c phenotype at 27°C of animals deficient in HID-1, a key component in the secretion of DCVs [43] (S6A Fig). HID-1 is expressed in all neuron and gut cells of *C. elegans*. Expression of HID-1 under the *rab-3* promoter in neurons in a *hid-1(sa722)* mutant background was sufficient to partially restore the rescuing effect of 2-AG on dauer formation (S6B Fig). In contrast, 2-AG failed to rescue dauer formation when HID-1 was expressed under the *ges-1* promoter in the gut of *hid-1* animals (S6B Fig). Taken together, these experiments indicate that diminished neural release of DCVs impairs the effect of 2-AG on cholesterol mobilization.

## Discussion

Although cannabis extracts have been used in folklore medicine for centuries, the past few years have increased interest in the medicinal use of cannabinoids, the bioactive components of the cannabis plant for treatment of many diseases of the nervous system. We have recently demonstrated that 2-AG and AEA, the best characterized eCBs reversed the blockade of intracellular trafficking of cholesterol in *C. elegans* [15]. Clearly, unraveling the molecular basis of cholesterol mobilization by eCBs in *C. elegans* could have important implications for a greater understanding of human pathological conditions associated with impaired cholesterol homeostasis. The endogenous cannabinoid 2-AG and AEA are synthesized within the brain and CNS [44]. Cannabinoids primarily activate G_αo_-coupled cannabinoid receptors 1 and 2 (CB1 and CB2)[20,22]. CB1 is localized primarily in the brain and CNS, whereas CB2 is restricted to the periphery and certain leukocytes [45]. Although initial reports suggested that nematodes lacked a canonical CB receptor [46,47], it has been determined that *C. elegans* also possesses cannabinoid-like receptors [17,18]. NPR-19 is a functional orthologue to the mammalian CB1/2, and NPR-32, a functional orthologue to GPR18 and GPR55 [17]. Here we show that the modulation of cholesterol homeostasis by 2-AG is independent of the GPCRs NPR-19 and NPR-32 signaling. Instead, we found that 2-AG displays interactions with components of the insulin/IGF1 signaling, a major endocrine pathway modulating DA production and dauer formation. We determined that reduction of the IIS pathway by mutations in the insulin/IGF receptor homolog *daf-2*, the phosphoinositide-3 kinase AGE-1/PI3K or the serine threonine kinase AKT-1 result in animals that are not rescued by 2-AG in a cholesterol depleted medium (Fig. 6). Since reduction of DAF-2/IIS pathway activity affects significantly 2-AG signaling, we asked whether insulin like peptide secretion control eCB-mediated cholesterol homeostasis. We found that the calcium-activated regulator of dense-core vesicle release (DCVs), UNC-31/CAPS, which functions in the nervous system to mediate release of insulin like peptides [38,40–42] is essential for 2-AG-mediated stimulation of cholesterol mobilization. This result combined with the requirement of *hid-1* for 2-AG signaling suggests that neural release of DCVs is required for 2-AG regulation of cholesterol homeostasis. We hypothesize that 2-AG stimulates cholesterol mobilization through the release of insulin-like peptides.

In agreement with early results [9] we have determined that after two generations of cholesterol depletion, DAF-16::GFP fusion protein exhibited significantly higher accumulation in the nucleus than did worms grown on cholesterol supplemented media (Fig 7A and S7 Fig). Exogenous 2-AG largely inhibited the nuclear localization of DAF-16 (Fig 7A), adding important evidence that the eCB stimulates the mobilization of cholesterol and its conversion into DA. Once synthesized, DA binds to DAF-12 and in target tissues liganded DAF-12 releases DAF-16 from the nucleus promoting reproductive growth and inhibiting dauer programs [9].

**Fig 7.**
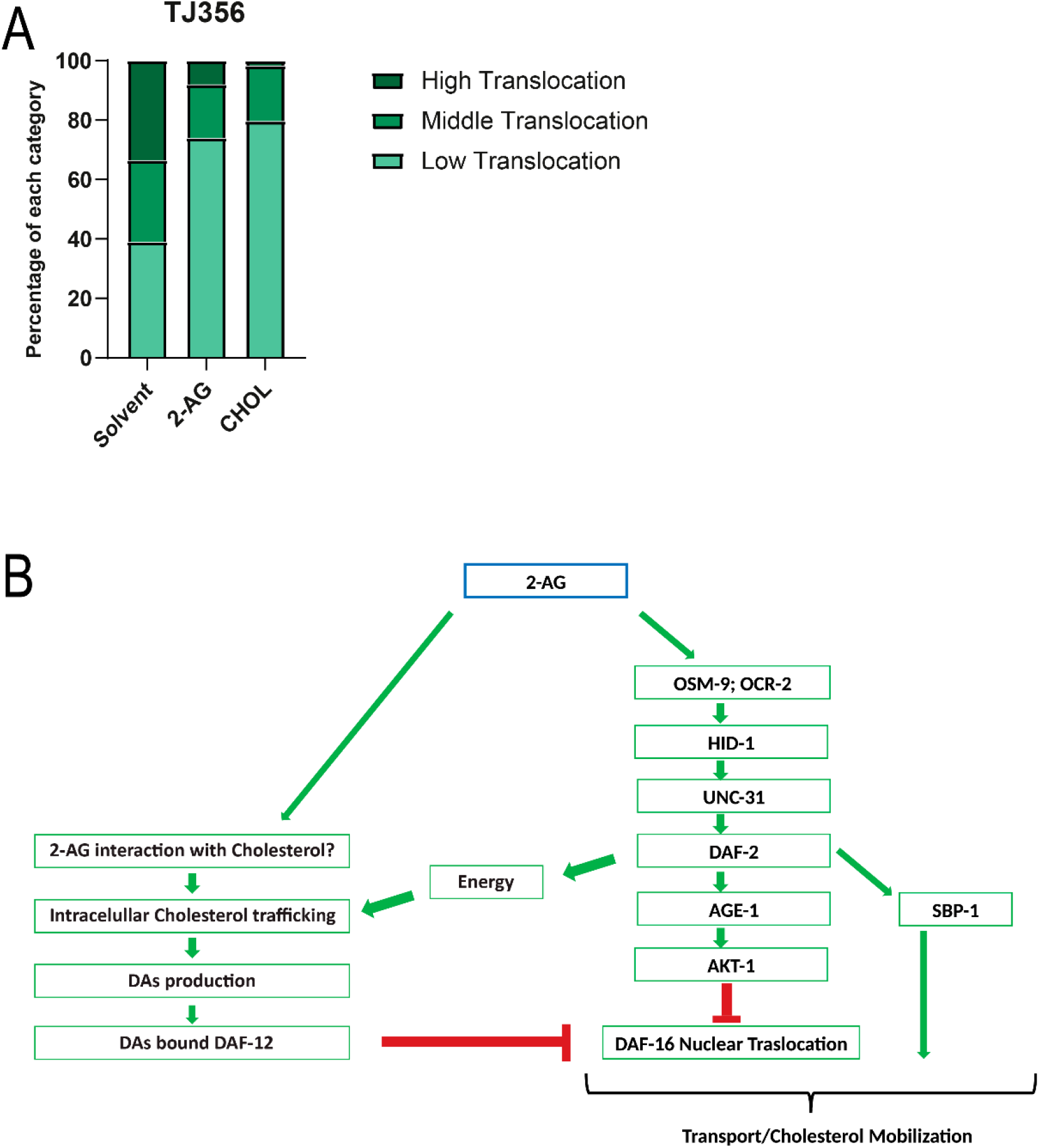
2-AG signaling inhibits nuclear translocation of DAF-16 induced by cholesterol depletion. (A) Nuclear translocation of DAF-16::GFP of worms grown under normal cholesterol concentration (CHOL) and after cholesterol depletion for two generations in the absence (solvent) or presence of 2-AG. 69 individual worms were analyzed for the solvent condition, 66 for 2-AG and 64 for CHOL. (B) Working model of 2-AG dependent mobilization of cholesterol. Under interaction of 2-AG with an unknown target in sensory neurons, multiple signaling mechanisms are likely to converge on activation of the IIS pathway and inhibition of DAF-16/FOXO. Signaling through the DAF-2 insulin receptor has several functions and could be involved in modulation of the intracellular traffic of cholesterol. The interaction of SBP-1 with the IIS pathway remains to be established, but is likely that such interaction is essential for 2-AG-dependent mobilization of cholesterol.

The simplest model consistent with previous [15] and present results is that 2-AG has a dual role, promoting the release of insulin peptides contained in DCVs and removing cholesterol from internal pools (Fig 7B). It is not known how and where worms store the internal sterol pools. Even though cholesterol is associated with cell membranes and interacts with multiple lipid species, very little is known about how lipids influence cholesterol trafficking. One of the few examples is the positive effect of the phospholipid lysobisphosphatidic acid on the trafficking of cholesterol through the endolysosomal compartment [48]. Owing to the huge diversity of membrane lipids, multiple other lipid species might emerge as additional modulators of the cholesterol trafficking process. eCBs are amphiphilic molecules derived from phospholipids that are unlikely to diffuse passively in the membrane. Several reports have shown that cholesterol behaves as a specific binding partner for eCBs [49,50]. Following an initial interaction of either 2-AG or AEA with cholesterol, mediated by the establishment of hydrogen bonds, they are attracted towards the membrane interior forming a molecular complex [49,50]. This raises the possibility that the interaction of 2-AG with cholesterol enhances the intracellular trafficking of sterols to steroidogenic tissues, positively affecting production of DAs. This regulated transport of cholesterol demands energy [51]. As upregulation of the insulin pathway is linked to increased metabolic rates [52], 2-AG-mediated activation of the DAF-2 pathway may induce a metabolic shift to provide the fuel needed to meet the high energy demands of *C. elegans* cholesterol mobilization (Fig 7B). In addition, stimulation of the IIS pathway should enforce, together with DA-bound DAF-12, the DAF-16/FOXO nuclear export to rescue the arrest of cholesterol depleted worms (Fig 7B).

Many of the physiological effects produced by eCBs are not completely understood. As we report here, some of them may reflect their influence on cholesterol trafficking. Interestingly, eCBs and eCB agonists are known to increase the hepatic expression of SREBP in mice [53]. Here we show that the simple orthologue of mammalian SREBP in *C. elegans*, SBP-1, plays an important role in 2-AG mobilization of cholesterol (Fig. 5B). Notably, RNAi of SBP-1 increases the penetrance of Daf-c defects seen in *daf-2 (e1370*) when grown in normal dietary cholesterol (S8 Fig). Although experimental evidence has demonstrated that *daf-2* controls the expression of numerous genes predicted to participate in fatty acid metabolism [54], it is not clear how DAF-2 and its regulatory targets regulates lipid metabolism. Our findings indicate that SBP-1 and IIS converge on a critical physiological process potentially related to the role of eCBs in cholesterol availability (Fig 7B). Future work should elucidate the nature of such interaction.

The specific molecular mechanisms through which *C. elegans* sensory neurons detect eCBs remain to be deciphered, however our findings suggest a key role for the *osm-9* and *ocr-2* TRPV genes in the control of cholesterol trafficking mediated by 2-AG. We found that both *osm-9* and *ocr-2* arrest already in the first generation without externally provided sterols (Fig 3), reminiscent of *fat-3* and *fat-4* mutants which are aberrant in cholesterol mobilization because they are unable to synthesize AEA and 2-AG [15]. Moreover, the arrest of *osm-9* and *ocr-2* mutants is not rescued by 2-AG, strongly suggesting that OSM-9/OCR-2 TRPV channel is essential for 2-AG-dependent cholesterol mobilization. Since the TRPV family encodes for non-selective cation channels with a preference for Ca^2+^ [30], it is possible that that OCR-2/OSM-9 may directly activate insulin secretion trough UNC-31, a Ca^2+^ dependent regulator of DCVs release.

Although the exact mode of activation of OSM-9 and OCR-2 in neurons is partly understood, our data are consistent with the hypothesis that 2-AG acts upstream of the TRPV channel. Interestingly, it has been suggested that G-protein-coupled lipid signaling pathways regulate TRPV channel signaling in chemosensory neurons [55]. This yet to be identified pathway could be specific inhibited by the cannabinoid based calcium blocker NMP331, that potently competes with 2-AG in cholesterol mobilization (S4B Fig).

Our insights in the role of eCBs in nematode cholesterol homeostasis has added an important new piece of information. Yet, the puzzle remains incomplete, and many mechanistic questions have yet to be answered. It seems plausible that both worms and mammals possess a fully functional eCB signaling pathway that regulates cholesterol homeostasis. Further dissecting cannabinoid regulation of lipid homeostasis in the context of larger endocrine networks should reveal how these processes alter disease states, health and possibly longevity.

## Materials and Methods

### Materials

2-AG and 2-AGE were purchased from Cayman Chemical (Ann Arbor, Michigan, USA) and stock solutions are in acetonitrile at 1 mg/ml and are stored at -80 ºC. Cholesterol, Dubelcco’s medium (DMEM) and antioxidant BHT were purchased from Sigma (Sigma-Aldrich, St. Louis, Missouri, USA). Δ4-DA and Δ7-DA were provided by Prof. H.-J. Knölker. All buffers, salts and chemical were reagent grade and were used without further treatments. Unless specified, all reagents were purchased from Merck or Sigma. The cannabinoid receptor ligands (NMP compounds) 241, 242, and 243 are named as compounds 40, 54 and 41 in reference 32. Compound NMP331 is named as compound 10 in reference 33.

### Nematode maintenance and strains

Standard *C. elegans* culture and molecular biology methods were used. Strains were cultured at 15°C or 20ºC on nematode growth media (NGM) agar plates with the *E. coli* OP 50 strain as a food source [56]. The wild type strain was Bristol N2. Some strains were provided by the *Caenorhabditis* Genetics Center (CGC) at the University of Minnesota. The strains *hid-1 (sa722), hid-1 (sa722); lin-15 (n765); jsEx897 [rab-3p-HID-1-GFP], hid-1 (sa722); lin-15 (n765); jsEx909 [ges-1p-HID-1-GFP], hid-1 (sa722); lin-15 (n765); jsEx896 [hid-1p-HID-1-GFP] were* kindly provided by D. Rayes. The strains used were: N2 Bristol (wild-type), *daf-7(e1372), daf-2(e1370), daf-2(e1368), fat-4 (ok958), ncr-1 (nr2022); ncr-2 (nr2023), npr-19 (ok2068), tyra-3(ok325), ckr-2(tm3082), gar-2(ok520), gar-3(vu78), gar-1(ok755), ser-2(ok2103), npr-11 (ok594), ser-1(ok345), ser-5(ok3087), npr-5(ok1583), dop-1(vs101), mod-1(ok103), octr-1(ok371), npr-16(ok1541), ser-7(tm1325), ser-4(ok512), dop-2(vs105), npr-24(ok312), dop-1(ok398), npr-32(ok2541), npr-35 (ok3258), gnrr-1 (ok238), age-1 (hx546), akt-1 (ok525), daf-4* (e1364), *daf-8* (e1393), *unc-64 (e246), unc-31(e928), daf-14 (m77), daf-28 (sa191), faah-4 (lt121), sbp-1 (ep79), nhr-8 (ok186), ocr-1 (ok132), ocr-2 (ok1711), ocr-3 (ok1559), ocr-4 (vs137), osm-9 (ok1677), daf-11 (m47), tph-1 (mg280), daf-36 (k114), cca-1 (ok3442*), *zIs356 [daf-16p::daf-16a/b::GFP + rol-6(su1006)]. hid-1 (sa722), hid-1 (sa722); lin-15 (n765); jsEx897 [rab-3p-HID-1-GFP], hid-1 (sa722); lin-15 (n765); jsEx909 [ges-1p-HID-1-GFP], hid-1 (sa722); lin-15 (n765); jsEx896 [hid-1p-HID-1-GFP]; xuEx1978 [Psra-6::Gcamp6(f), Psra-6::DsRed]*.

### Preparation of sterol-depleted plates and sterol-deprived worm culture

To obtain sterol-free conditions, agar was replaced by ultrapure agarose and peptone was omitted from plates as described earlier [15]. Briefly, agarose was washed three times overnight with chloroform to deplete the trace sterols in it. Salt composition was kept identical to NGM plates. As a food source, *E. coli* NA22 grown overnight in sterol-free culture medium DMEM was used. Bacteria were rinsed with M9 buffer and 20 times concentrated. Bleached embryos were grown for one generation on sterol-free agarose plates. The resulting gravid adults were bleached and the obtained embryos were used in various assays.

### Generation of *daf-7; npr-19* and *npr-19;npr-32* double mutants

The *daf-7* and *npr-32* mutants were each crossed into *npr-19* to generate double mutants by standard methods. Crosses were confirmed by PCR genotyping and constitutive dauer arrest at 25ºC.

### Dauer formation assays

Dauer assays were performed as previously described [15]. In general, 60–80 L1s or embryos were transferred to NGM plates seeded with *E. coli* (HT115 or OP50). eCBs (final concentration 50 μM) or NMP331 (final concentration 1 μM or as indicated in Supplementary Fig. 1) were added to the bacteria immediately prior to seeding. The final concentrations of these compounds were calculated according to the volume of the NGM agar used for the preparation of the plates. Depending on the sort of essays after 3, 4 or 5 days the dauer percentage was scored. Δ7-DA was used alternatively in dauer rescue experiments at a concentration of 90nM.

### Lipid Extraction and endocannabinoid analysis by HPLC-MS/MS

The protocol is adaptation of Folch (1957) [57]. Briefly, lipid extracts were made from approximately 200 mg of frozen worm pellets grown at 20 °C. Pellets were washed with M9 buffer, then re-suspended in 1.3 ml pure methanol and sonicated three times for 30 seconds. After sonication 2.6 ml of chloroform and 1.3 ml 0.5 M KCl/0.08 M H3PO4 were added. Butylated hydroxytoluene (BHT, 0.005 % v/v) was added to prevent lipid oxidation. Samples were then sonicated for 15 min, vortexed twice for 1 min and centrifuged for 10 min at 2.000 x g to induce phase separation. The lower, hydrophobic phase was collected, dried under constant nitrogen stream, re-suspended in 100 μl of acetonitrile and loaded into dark caramel tubes using a glass pipette. 2-AG was quantified from nematode samples by liquid chromatography (Ultimate 3000 RSLC Dionex, Thermo Scientific) coupled with an ESI triple quadrupole mass spectrometer (TSQ Quantum Access Max (QQQ), Thermo-Scientific) as previously described [15].

### DAF-16::GFP expression analysis

An stably integrated DAF-16::GFP, TJ356 was used. The expression of GFP was observed using a Nikon Eclipse 800 microscope equipped with a fluorescent light source. The images were captured with a Andor Clara digital camera.

To observe GFP distribution, 15-25 animals were mounted to an agar pad containing 20 mM sodium azide and analyzed immediately. No animal on the pad for more 10 min was scored. DAF-16::GFP distribution was categorized based on the accumulation of GFP in the nucleus or the cytoplasm (diffuse form) in the whole animal. Individual animals were classified based on the presence of nuclear DAF-16::GFP in approximately 90%, 70% (High translocation), 50%, 30% (Middle traslocation), 10% or none (Low Translocation) of the body cells, as shown in Figure Supplementary 7. All the animals scored were at the stage of L2 after 72 hs of growing in the second generation under free sterol conditions.

### Two-electrode voltage clamp in *Xenopus oocytes*

For electrophysiological recording in *Xenopus levis* oocytes, OSM-9, OCR-2 subunits subcloned into a modified pGEMHE vector were used. cRNAs were in vitro transcribed from linearized plasmid DNA templates using RiboMAXTM Large Scale RNA Production System (Promega, Madison, WI, USA). Xenopus oocytes were injected with 50 nl of RNase-free water containing 1.0 ng of cRNA (at a 1:1 molar ratio for heteromeric receptors) and maintained in Barth’s solution [in mM: NaCl 88, Ca(NO3)2 0.33, CaCl2 0.41, KCl 1, MgSO4 0.82, NaHCO3 2.4, HEPES 10] at 18°C. Electrophysiological recordings were performed at −60 mV under two-electrode voltage-clamp with an Oocyte Clamp OC-725B or C amplifier (Warner Instruments Corporation, Hamden, CT, USA). Recordings were filtered at a corner frequency of 10 Hz using a 900 BT Tunable Active

Filter (Frequency Devices Inc., Ottawa, IL, USA). Data acquisition was performed using a Patch Panel PP-50 LAB/1 interphase (Warner Instruments Corp., Hamden, CT, USA) at a rate of 10 points per second. Both voltage and current electrodes were filled with 3M KCl and had resistances of ∼1MΩ. Data were analysed using Clampfit from the pClamp 6.1 software (Molecular Devices, Sunnyvale, CA). During electrophysiological recordings 100 μM 2-AG was added to the perfusion solution. Recording was performed at room temperature and heat-stimulation (∼ 36 °C) by perfusion of heated Barth’s. The temperature of perfused bath solutions was checked with a TC-344B temperature controller (Warner Instruments) located near the oocytes. Mean ± SEM of current amplitudes of responses to temperature, 100 μM 2-AG and temperature plus 100 μM 2-AG in oocytes injected with either OSM-9, OCR-2 or OSM-9/OCR-2 were calculated using Prism 6 software (GraphPad Software Inc., La Jolla, CA, USA).

### ASH calcium imaging

Animals expressing GCaMP6 in the ASH (xuEx1978 [Psra-6::Gcamp6(f), Psra-6::DsRed]) were immobilized in a PDMS microfluidic olfactory chip [58,59] and exposed to either a control buffer or a stimulus buffer containing 100mM 2-AG (Cayman Chemical Co. #62160). The required volume of 2-AG required to make 100mM solution was dissolved in 0.1% ethanol before being added to S-Basal buffer. Since the 2-AG stock solution was dissolved in acetonitrile, we added an equivalent volume of acetonitrile and 0.1% ethanol S-basal to make the control buffer. This ensured the responses were specific to 2-AG. The stimulus protocol was based on previously described exposure experiments [55]. Animals were allowed to acclimate in the olfactory chip for at least 5 minutes before recording. Recording was performed at 10x magnification on an AxioObserver A1 inverted microscope (Zeiss) connected to a Sola SE Light Engine (Lumencor) and an ORCA-Flash 4.0 digital CMOS camera (Hamamatsu). Micromanager Software [60] was used to control image acquisition. Recording was performed at 10 frames per second and 4×4 image binning. An Arduino was used to control pinch valves to direct stimulus and control buffer to the nose of animals in the following sequence: 6 second baseline, 4 second stimulation, 10 second interstimulus interval, 4 second stimulation, 6 second washout. GCaMP fluorescence was extracted using Matlab (Mathworks) scripts and the results were plotted using GraphPad Prism.

## Acknowledgments

Supported by grants of Richard Lounsbery Foundation (to D.d.M) and NIH GM1140480 (to M.J.A). Some strains were provided by the CGC, which is funded by NIH Office of Research Infrastructure Programs (P40 OD010440). We thank Sider Penkov and Teymuras Kurzchalia for sharing unpublished data, to Luisa Cochella, Cecilia Mansilla and Claudia Banchio for critically reading the manuscript and to Diego Rayes and Shawn Xu for the worm strains.

## Supporting information captions

**S1 Fig. TPH-1 is not required for 2-AG-dependent cholesterol mobilization**. (A) N2 and *tph-1* were grown for two generations in media with 0 μg/ml cholesterol at 20ºC. Mann-Whitney rank sum test, *p < 0.001. t-test, **p < 0.001. All values from n = 3 independent experiments show as Mean ± SEM. N2 is the *C. elegans* wild-type strain.

**S2 Fig. 2-AG did not elicit current in Xenopus oocytes expressing OSM-9 and OCR-2 channels**. (A) Representative traces of responses (above) and temperature (below) in *Xenopus oocytes* injected with cRNAs encoding OSM-9, OCR-2, OSM-9/OCR-2. (B) Mean ± SEM of heat evoked currents. Amplitudes were calculated by measuring the differences between the peak inward currents and baseline marked with dotted lines (***p = 0.0001, n ≥ 5 per group, ANOVA followed by a Bonferroni’s multi-comparison test). (C) Representative traces of responses to 100μM 2-AG either at room temperature (grey traces) or during a temperature ramp in oocytes expressing OSM-9, OCR-2 or OSM-9/OCR-2. (D) Mean ± SEM of current amplitudes of responses to temperature, 100 μM 2-AG and temperature plus 100 μM 2-AG in oocytes injected with either OSM-9, OCR-2 or OSM-9/OCR-2 (**** p < 0.0001, n ≥ 3 oocytes per group, two-way ANOVA followed by a Bonferroni multi-comparison test).

**S3 Fig. ASH calcium levels are not affected by 2-AG**. Calcium imaging traces of animals expressing GCaMP6 in the ASH (xuEx1978 [Psra-6::Gcamp6(f), Psra-6::DsRed]) during exposure to 2-AG. Average responses for all animals (A) are indicated by the dark blue line and the shaded area represents SEM. The same results are shown as individual traces for each animal (B). The dark grey bars indicate 4 second exposure to buffer containing 100mM 2-AG.

**S4 Fig. NMP331 induce a dauer-like formation in N2 starved for sterols and enhances dauer formation in *daf-7* mutants which can be overcome by 2-AG supplementation**. (A) N2 exhibit a dauer-like phenotype when is exposed to 1 μM NMP31 in a sterol-free medium at 25 ºC during one generation. The black straight line represents 0.25mm. (B) 2-AG antagonizes the effect of 1 μM of NMP331 in a *daf-7* worm at 20 ºC. All pairwise multiple comparison procedures (Holm-Sidak method), *p < 0.001. All values from n = 3 independent experiments show as Mean ± SEM.

**S5 Fig. Mobilization of cholesterol by 2-AG is independent of the *daf-7* pathway**. (A). *Daf-4* was grown at 20ºC while *daf-14* and *daf-8* were grown at 25 ºC under normal dietary cholesterol (13 μM). t-test, *p < 0.001. **p < 0.001. ***p < 0.002. All values from n = 3 independent experiments show as Mean ± SEM. (B) NMP331 does not enhances the daf-c phenotype of *daf-2* mutants. Mann-Whitney rank sum test, *p < 0.001. All values from n ≥ 3 independent experiments show as Mean ± SEM. ns = not significant.

**S6 Fig. Mobilization of cholesterol by 2-AG is dependent of *hid-1***. (A) *Hid-1* is required for 2-AG dependent mobilization of cholesterol. Worms were grown at 27 ºC under normal dietary cholesterol. All pairwise multiple comparison procedures (Holm-Sidak method), *p < 0.05. All values from n = 3 independent experiments show as Mean ± SEM. ns = not significant. (B) HID-1 expression in the neurons in a *hid-1* background restores the 2-AG-dependent mobilization of cholesterol. Worms were grown at 27 ºC under normal dietary cholesterol. Mann-Whitney rank sum test, *p < 0.001. All values from n ≥ 3 independent experiments show as Mean ± SEM. ns = not significant.

**S7 Fig. DAF-16::GFP nuclear translocation groups classification**. Translocation categories were placed into 3 classes as the picture shows. Animals which exhibited DAF-16::GFP nuclear accumulation between 0-10% of total DAF-16::GFP were classified as Low Translocation (LowT), 30-50%, as Middle Translocation (MiddleT) and 70-90%, High Translocation (HighT). All animals shown are in a L2 larva stage.

**S8 Fig. sbp-1 RNAi increases the *daf-c* phenotype of *daf-2***. *daf-2* was grown in 13 μM cholesterol at 20ºC. Mann-Whitney rank sum test, *p < 0.005. All values from n = 3 independent experiments show as Mean ± SEM.

**S1 Table. Orthologues of CB1/2 grown for two generations in the absence of cholesterol arrest as L2-like larvae**.

**S2 Table. RNAi enhancer screen on *daf-7 (e1372)* worms**. For details see text.

